# Cryptosporulation in *Kurthia* spp. forces a rethinking of asporogenesis in Firmicutes

**DOI:** 10.1101/2020.06.18.160689

**Authors:** Sevasti Filippidou, Mathilda Fatton, Thomas Junier, Matthieu Berge, Daniel Poppleton, Thorsten B Blum, Marek Kaminek, Adolfo Odriozola, Jochen Blom, Shannon L Johnson, Jan Pieter Abrahams, Patrick S Chain, Simonetta Gribaldo, Elitza I. Tocheva, Benoît Zuber, Patrick H Viollier, Pilar Junier

## Abstract

Sporulation is a complex morphophysiological process resulting in a cellular structure that is more resistant than the vegetative form. In Firmicutes, this structure is produced within the mother cell, and is called an endospore. Endospore formation is thought to have evolved in the common ancestor of Firmicutes. However, sporulation has apparently been lost in some extant lineages that are defined as *asporogenic*. We isolated strain 11kri321, a representative of the genus *Kurthia*, from an oligotrophic geothermal reservoir. While *Kurthia* spp. is considered to comprise only asporogenic species, strain 11kri321 produced spores. Genomic reconstruction of the sporulation pathway shows elements typical of sporulation in Bacilli, including the signaling for sporulation onset. However, key genes were missing, including those involved in engulfment and dipicolinic acid synthesis. Based on the results for strain 11kri321, sporulation was investigated in other *Kurthia* strains. Genes involved in signaling, cell division and spore coat formation were detected in three available *Kurthia* genomes. Moreover, endosporulation was clearly visualized in at least two of the four strains tested. These results show that *Kurthia* is an endospore-forming Firmicute lineage. However, the genetic background of sporulation in this genus deviates from the known sporulation pathway in Firmicutes and even within Bacilli, suggesting that a revision of the minimal set of genes used for genomic detection of sporulation is required. Based on our findings we propose the term *cryptosporulant* to refer to putative asporogenic Firmicutes for which a detailed genomic and physiological characterization of sporulating is missing.

**Importance:** Endospore-forming Firmicutes include many environmental and medical relevant bacterial clades. In these microorganisms, the ability to produce endospores is essential for survival in the environment and even for pathogenesis. The minimum core of genes required to produce a viable and resistant spore, the distinction between endospore-forming and asporogenic groups, as well as the evolution of sporulation have been a subject of investigation and debate for decades. Here, we demonstrate endosporulation in the genus *Kurthia,* considered as asporogenic. Morphological, physiological and genomic analyses were undertaken to demonstrate that sporulation is not lost within this lineage. Based on our results we propose a re-examination of the minimal genetic requirements of sporulation and the use of the term *cryptosporulant* to describe lineages of Firmicutes that have not previously been observed to sporulate, but for which a detailed analysis is still missing.

## Introduction

Sporulation is a morphophysiological response to unfavorable environmental conditions involving a sophisticated genetic mechanism of cellular division and differentiation. In Firmicutes, the process of sporulation is called endosporulation, because it is the result of an asymmetrical cell division that leads to the formation of a mature spore within a mother cell (1). Endosporulation is thought to have emerged in the last common ancestor of Firmicutes (2) but appears to be have been lost or become inactive in many extant descendants (3). When sporulation is no longer possible, and despite the presence of some genomic remnants of sporulation, the phenotype of the organism is defined as *asporogenic*. The advent of genome sequencing technologies has opened the door to re-investigate supposed asporogenic species. For instance, contrary to the previous physiological knowledge, the analysis of the genomes of *Carboxydothermus hydrogenoformans* and *Ruminococcus bromii* suggested that these species could produce spores. This prompted researchers to experimentally demonstrate sporulation in both groups (4, 5).

A series of approximately 60 genes are proposed to encode the minimal genetic core for sporulation. This minimal set has been defined based either on comparative genomics of known endospore-forming species (6), or alternatively, on the set of genes derived from the genomic analysis of specific species (for instance, analysis tools such as Genome Properties (7) uses the sporulation genes found in *C. hydrogenoformans*). The idea of a minimal set of sporulation-related genes has been crucial in the investigation of unusual endospore-formers such as segmented filamentous bacteria, which despite their small genomes have all the predicted set of core sporulation genes (8, 9). This defined set of genes is also relevant when judging the claims of sporulation in supposedly asporogenic clades outside Firmicutes, such as in the case of reports of endospore formation in *Mycobacterium* spp. (10, 11).

Although the mechanisms underlying the maintenance of endospore formation are still poorly understood, one can hypothesize that given its high energetic cost and genetic complexity, bacteria might lose the ability to produce spores under constant favorable conditions for vegetative growth. This has been shown experimentally in model endospore-forming species in which an asporogenic phenotype is the result of the inactivation or loss of a considerable fraction of sporulation genes (12). However, under highly variable environmental conditions, sporulation is considered to be a beneficial trait that improves survival and dispersal (13). Accordingly, a recent culturing effort combined with extensive genomic analysis of the human microbiota has shown that sporulation is widely spread in bacteria inhabiting the human gut, an environment naturally subjected to high physicochemical fluctuation (14).

Ecosystems with extreme environmental conditions, such as geothermal sites, also harbor a larger diversity of endospore-formers (15). Therefore, in our quest to better characterize the diversity of environmental endospore-forming Firmicutes, multiple enrichments were conducted from samples collected in geothermal environments. The resulting strains were screened for their ability to form spores, without any prior bias regarding their phylogenetic affiliation. In this way, strain 11kri321 was isolated from the geothermal spring of Krinides, Kavala, Greece. The strain was found to belong to the genus *Kurthia*, which is classified as an asporogenic firmicute. In this study we present the physiological and genomic evidence demonstrating sporulation in *Kurthia*. Based on our findings, we discuss the implications of defining a truly asporogenic lifestyle within Firmicutes.

## Results

### Characterization of the isolate

Strain 11kri321 was isolated from a biofilm and was characterized as a Gram-positive bacterium (Fig. 1A). Affiliation of the strain to the genus *Kurthia* was supported by 16S rDNA sequencing and average amino acid identity (AAI) analysis of its full genome. The AAI values for the comparison between strain 11kri321 are 68.88% with *Kurthia massiliensis*, 68.50% with *Kurthia huakuii* and 68.58% with *Kurthia senegalensis*, which are all above the suggested threshold of 60% used for the definition of a bacterial genus (16). The phylogenetic placement of strain 11kri321 was further verified by a phylogenomic analysis showing the phylogenetic position of *Kurthia* relative to other members of the Firmicutes (Fig. 1B).

**Figure 1.**
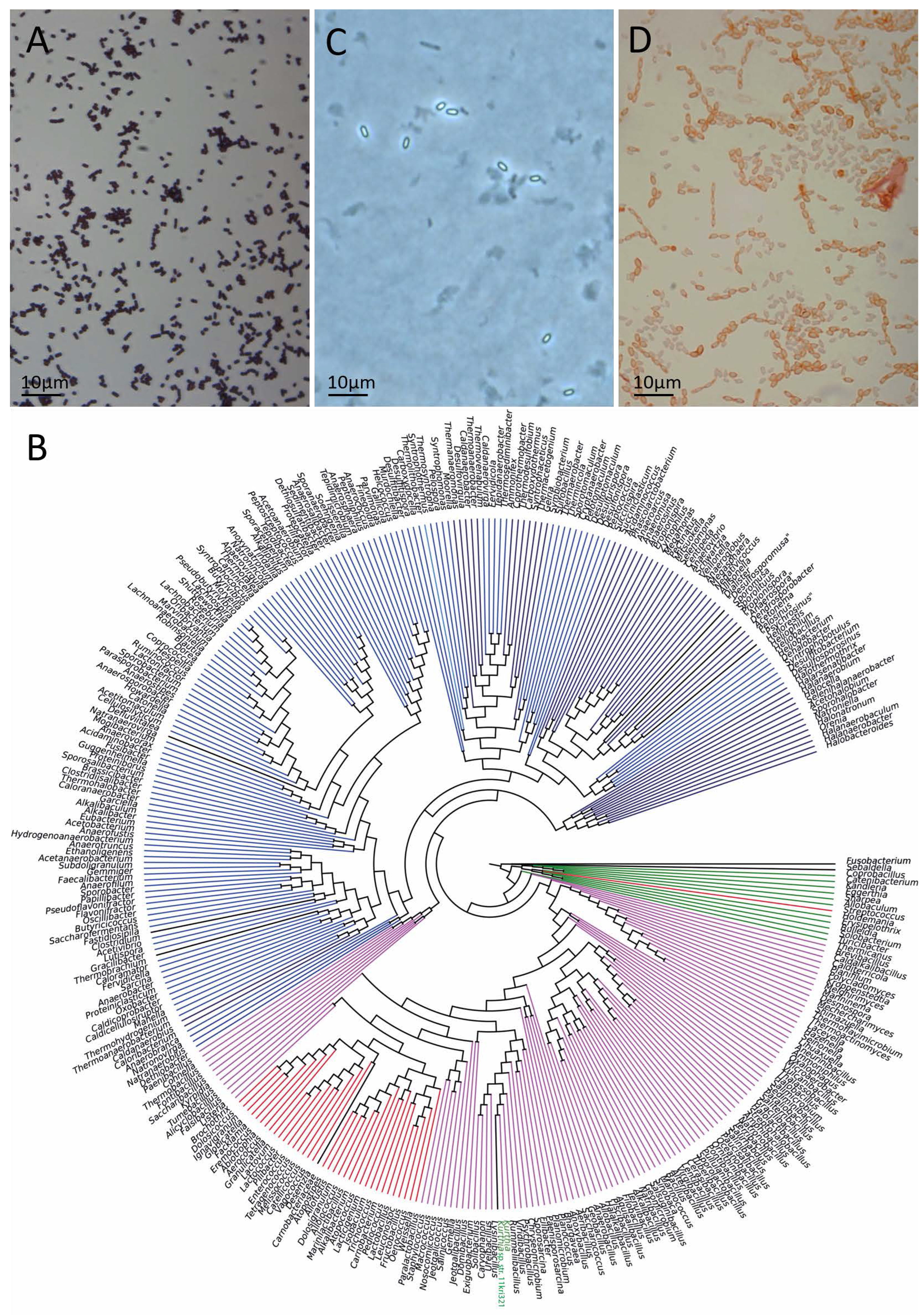
Morphological characterization and phylogenetic placement of *Kurthia* sp. str. 11kri32. Strain 11kri321 is a Gram-positive bacterium **(A)**. A Maximum-likelihood tree shows the positioning of the *Kurthia* genus among endospore-forming Firmicutes genera **(B)**. Branches are colored by class (magenta: Bacillales; blue: Clostridiales; green: Erysipelotrichales; navy: Halanaerobiales; crimson: Lactobacillales; mediumblue: Natranaerobiales; darkblue: Selenomonadales; midnightblue: Thermoanaerobacterales; dodgerblue: Thermolithobacterales; *Kurthia* genus, along with *Kurthia* sp. str. 11kri321, is highlighted in green lettering. Spore-like structures appeared phase-bright **(C)** and retained the malachite green staining **(D)**, as endospores of *Bacillus subtilis*.

In addition, we performed a physiological characterization of the strain. Strain 11kri321 grew at a pH between 5.5 and 11.5. Its optimal growth temperature was 25°C, however it could grow at temperatures between 20 and 45°C. The *in situ* pH of the site was 9 and the temperature was 29°C, both of which are within the limits of tolerance for all previously described *Kurthia* spp. (17), and within the values established for the vegetative growth of strain 11kri321. All these observations suggest that strain 11kri321 could have survived in a vegetative state in the borehole.

Although the genus *Kurthia* is considered asporogenic (18), spore-like structures from a nutrient-deprived culture of *Kurthia* sp. str. 11kri321 refracted light under the light microscope and appeared phase-bright (Fig. 1C), suggesting that nutrient starvation can, *in vitro*, induce spore-formation in this strain. In order to better characterize these structures, we performed staining with malachite green (Schaeffer-Foulton stain). This chemical was retained by the *Kurthia* spore-like structures but not by vegetative cells (Fig. 1D).

A more detailed morphological characterization of these spore-like structures was conducted using cryo electron tomography (cryo-ET). The average spore-like structure measured 700×1200 nm. Each spore-like structure was composed of features unique to endospores (19, 20) such as a core, thick cortex, inner and outer spore membranes (IsM and OsM, respectively), and a spore coat (Fig. 2A-B). Due to the thickness of the spores, the bilayer lipid of the IsM and OsM could not be resolved. Filamentous proteinaceous appendages analogous to an exosporium emerged from both spore poles and connected to the coat (Fig. 2A-B).

**Figure 2.**
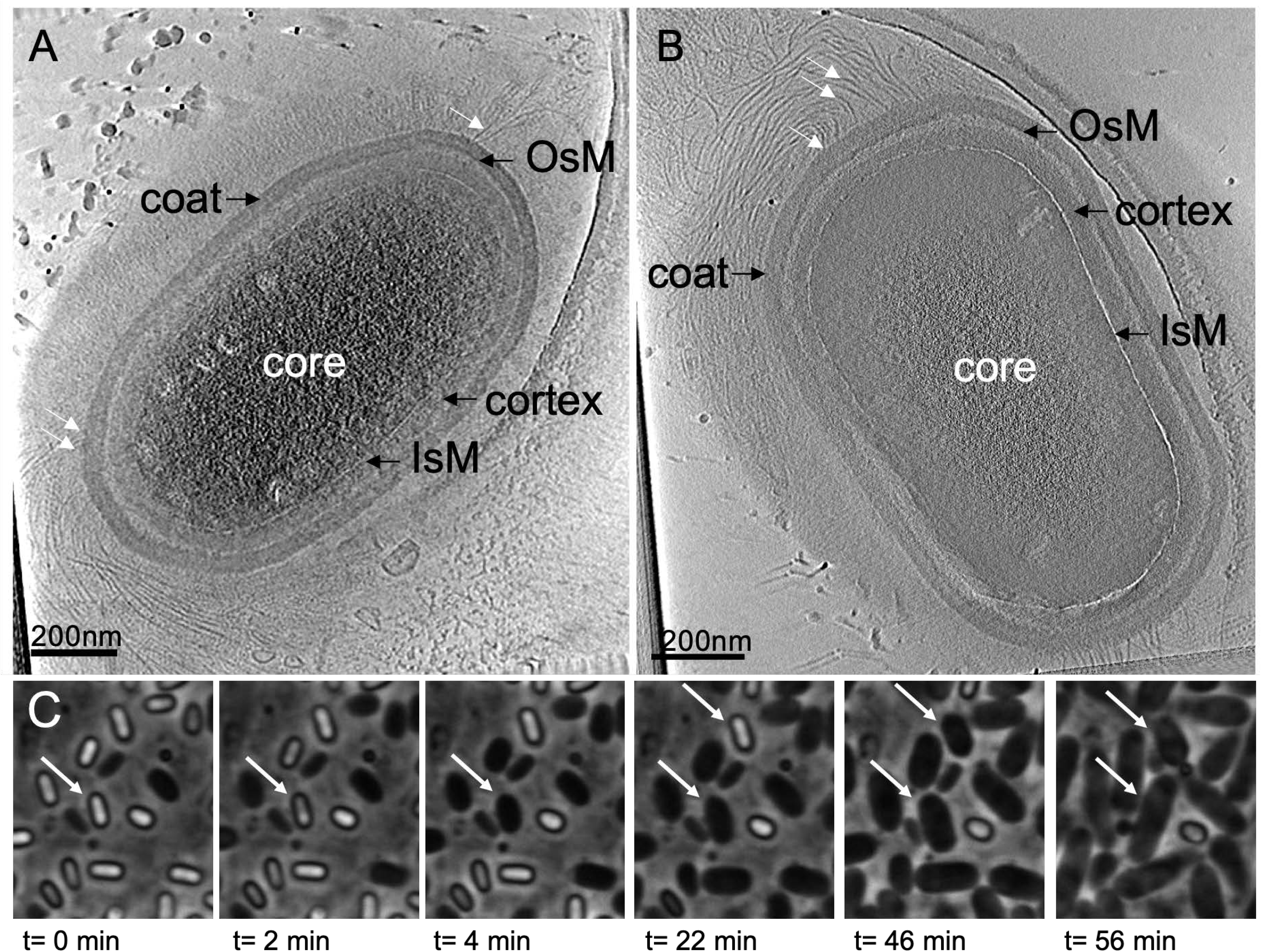
Detailed morphological characterization of the spore-like structures of *Kurthia* sp. str. 11kri321 by cryo electron tomography (cryo-ET) and monitoring of their germination by light microscopy. **(A-B)** Cryo-ET images of the spores of *Kurthia* sp. str. 11kri321, show the inner (IsM) and outer (OsM) spore membranes, spore cortex, spore coat, and filamentous appendages (white arrows). (**C)** Pictures from snapshots of the time-lapse movie in Supplementary Movie 1 showing key stages in the germination of *Kurthia* sp. str. 11kri321 spores.

To confirm that the observed phase-bright bodies were indeed viable spores, germination of *Kurthia* sp. str. 11kri321 was monitored by time-lapse microscopy (Fig. 2C; Supplementary Movie 1). We observed a shift in bright phase, swelling of the cells, and elongation, resulting in the characteristic short-rod-shaped of dividing vegetative cells. Only 6% of the spores did not germinate, a rate comparable to *B. subtilis* (16).

### Genomic imprints of sporulation in *Kurthia* sp. strain 11kri321

The 2.9 Mbp genome of *Kurthia* sp. str. 11kri321 was screened for 213 sporulation gene homologs, from 34 endospore-forming Firmicutes (Supplementary Table 1). In total, 43 sporulation genes were detected and assigned to the different stages of the endospore-formation pathway (Fig. 3A). Genes homologs to *sigH* and *spo0A*, encoding two of the main regulators responsible for controlling the onset of sporulation (18, 21), were found in the genome. Concerning the activation of these regulators, a homolog to *spo0F* (Spo0F is the response regulator in Bacilli (22)), was identified in the genome of *Kurthia* sp. str. 11kri321. The s*po0B* gene was not detected; however, the *obg* gene, which encodes for a Spo0B-associated GTP-binding protein, was found. In fact, by lowering the similarity threshold of detection of sporulation-related genes, we detected several proteins with conserved Spo0B domains, but those need to be validated experimentally. Moreover, a gene encoding for the protein KinA (activator of Spo0A (23)) was also detected in the genome of *Kurthia* sp. str. 11kri321.

**Figure 3.**
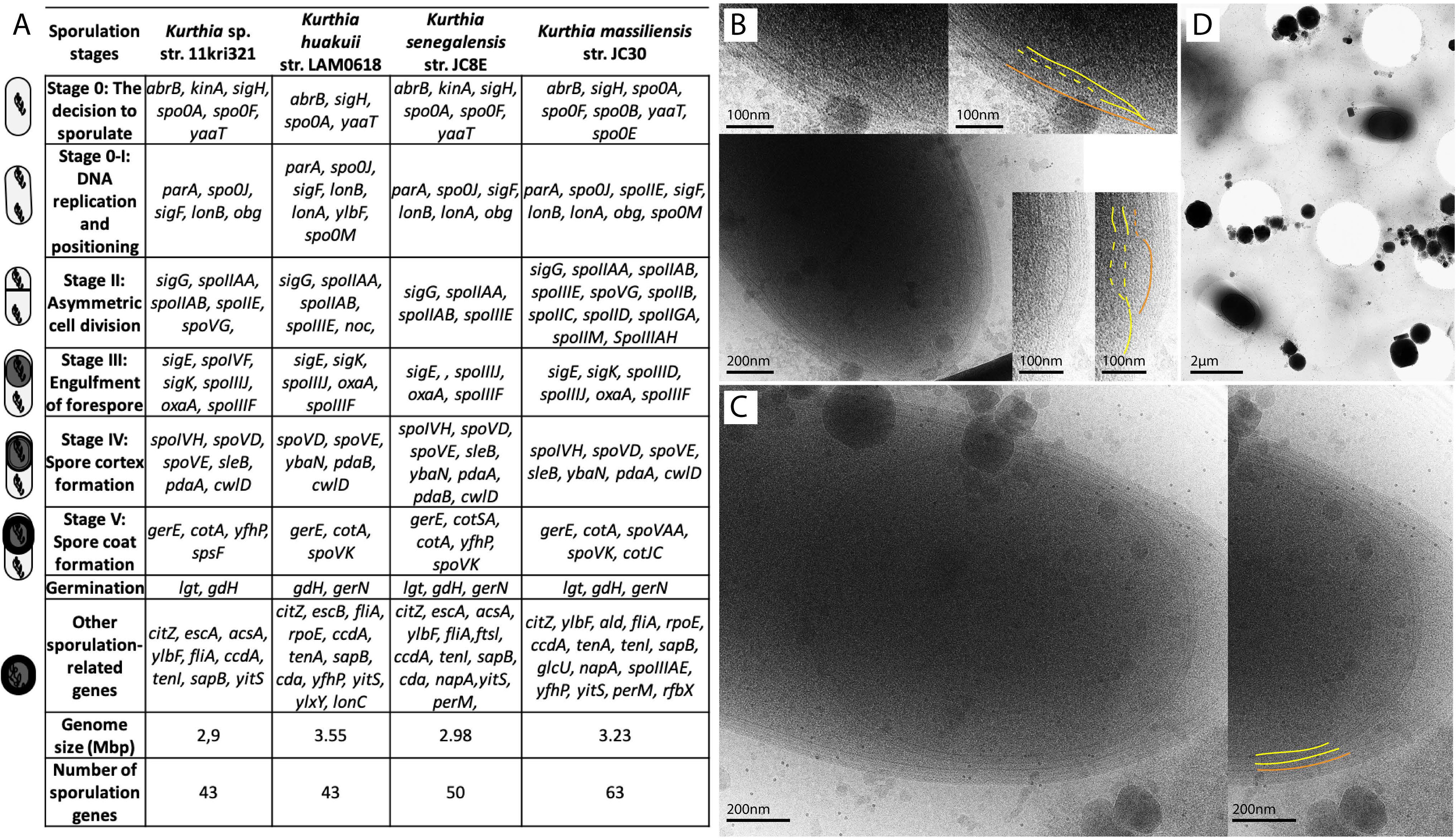
Comparative analysis of sporulation genes and 2-D cryo electron microscopy (cryo-EM) imaging of engulfment in *Kurthia* spp. **(A)** The sporulation genes found in the four genomes of *Kurthia* spp. are presented in the order of the sporulation stage in which they are expressed. Genes expressed in multiple stages are presented only once in this table. **(B)** Cryo-EM projection images reveal the steps of engulfment in *K. huakuii* str. LAM0618. The areas in which engulfment leads to the production of a pre-spore with two membranes (yellow) by the engulfment of the membrane from the mother cell (orange) are shown enlarged in the inserts. **(C)** A projection image of *K. huakuii* str. LAM0618 at the end of the engulfment step with the mature endospore (two membranes highlighted in yellow) inside the mother cell (inner membrane highlighted in orange). **(D)** A projection image showing endospores inside two mother cells of *K. senegalensis* str. JC8E.

Genes encoding four highly conserved (24) sigma factors (SigF, SigE, SigG and SigK) were found in the genome of *Kurthia* sp. str. 11kri321. In addition to these sigma factors, homologs encoding two of their known regulators, SpoIIAA and SpoIIAB (25, 26), were also detected. On the contrary, the *spoIIIAABCDEFGH* operon, responsible to regulate expression of the sigma factors (25, 26) involved in sporulation, was missing.

Another key stage of endospore-formation is the packaging of the entire genome within the forespore (27). A homolog to *spoIIE*, encoding the DNA transport protein SpoIIIE that carries out the translocation of the remaining fraction of the genome in Bacilli and Clostridia, was also identified in the genome of *Kurthia* sp. str. 11kri321. None of the genes known to be involved in the zipping process, postulated during engulfment in *Bacillus*, were identified. Although the sporulation-specific sigma G factor was found in *Kurthia* sp. str. 11kri321, genes coding for proteins with homology to known small acid-soluble proteins (SASPs) could not be identified. Finally, among the genes believed to be involved in the formation of the spore cortex in *B. subtilis,* only *spoVD* and *spoVE* were identified in the genome of *Kurthia* sp. str. 11kri321. Likewise, a much smaller set of genes encoding proteins clearly related to the formation of the spore coat (only *cotA*), were present.

### Presence of dipicolinic acid in the spores of strain 11kri321

An important molecule produced in the mother cell of endospore-forming Firmicutes and introduced to the spore at later stages of sporulation is dipicolinic acid (DPA). DPA is responsible for accumulation of minerals (especially calcium ions) in the spore core, to create a more stable core and to guarantee resistance to wet heat (28). The two genes that encode the DPA synthetase subunits A and B (*spoVFA*, *spoVFB*) were not detected in the genome of *Kurthia* sp. str. 11kri321 and accordingly, we were unable to detect any DPA from spore preparations (Supplementary Fig. 1). *Kurthia* sp. str. 11kri321 also lacks the gene *ger3*, as it is the case of *B. subtilis* mutants able to produce stable DPA-free spores (29, 30).

### Sporulation in other *Kurthia* species

In order to determine whether sporulation may be characteristic of the entire genus, we attempted chemical induction of sporulation in three additional *Kurthia* species (*K. huakuii* str. LAM0618, *K. senegalensis* str. JC8E, and *K. massiliensis* str. JC30), alongside strain 11kri321. All were tested for induced sporulation using 10% glycerol or nutrient deprivation in two sporulation media, with and without a carbon source (31, 32). All *Kurthia* strains were able to produce phase-bright bodies on most sporulation media, albeit at a frequency never exceeding 10% (Supplementary Fig. 2). The draft genomes of these *Kurthia* strains were also available and were screened for spore-related genes. A total of 43, 50, and 63 sporulation genes were found in *K. huakuii* str. LAM0618, *K. senegalensis* str. JC8E and *K. massiliensis* str. JC30 genomes, respectively (Fig. 3A).

Among the genes with a clear function, a striking difference between the genes identified in *K. massiliensis* and the other strains of *Kurthia* is found at the stage of decision-making to commit to sporulation. While homologs to SigH, Spo0A and Spo0F were found in all *Kurthia* genomes, Spo0B, which has a role in signal transduction on the *yaaT* gene, was only found in *K. massiliensis*. The *spoIIIAH* gene was detected in the genome of *K. massiliensis* but was absent from the other screened genomes. The other sporulation genes with a known function appear to be conserved within *Kurthia* spp. A comparative representation of the four *Kurthia* genomes to *Bacillus subtilis* subsp. *subtilis* str. 168 is shown in Figure 4, including potential sporulation operons deduced from the genomic analysis.

**Figure 4.**
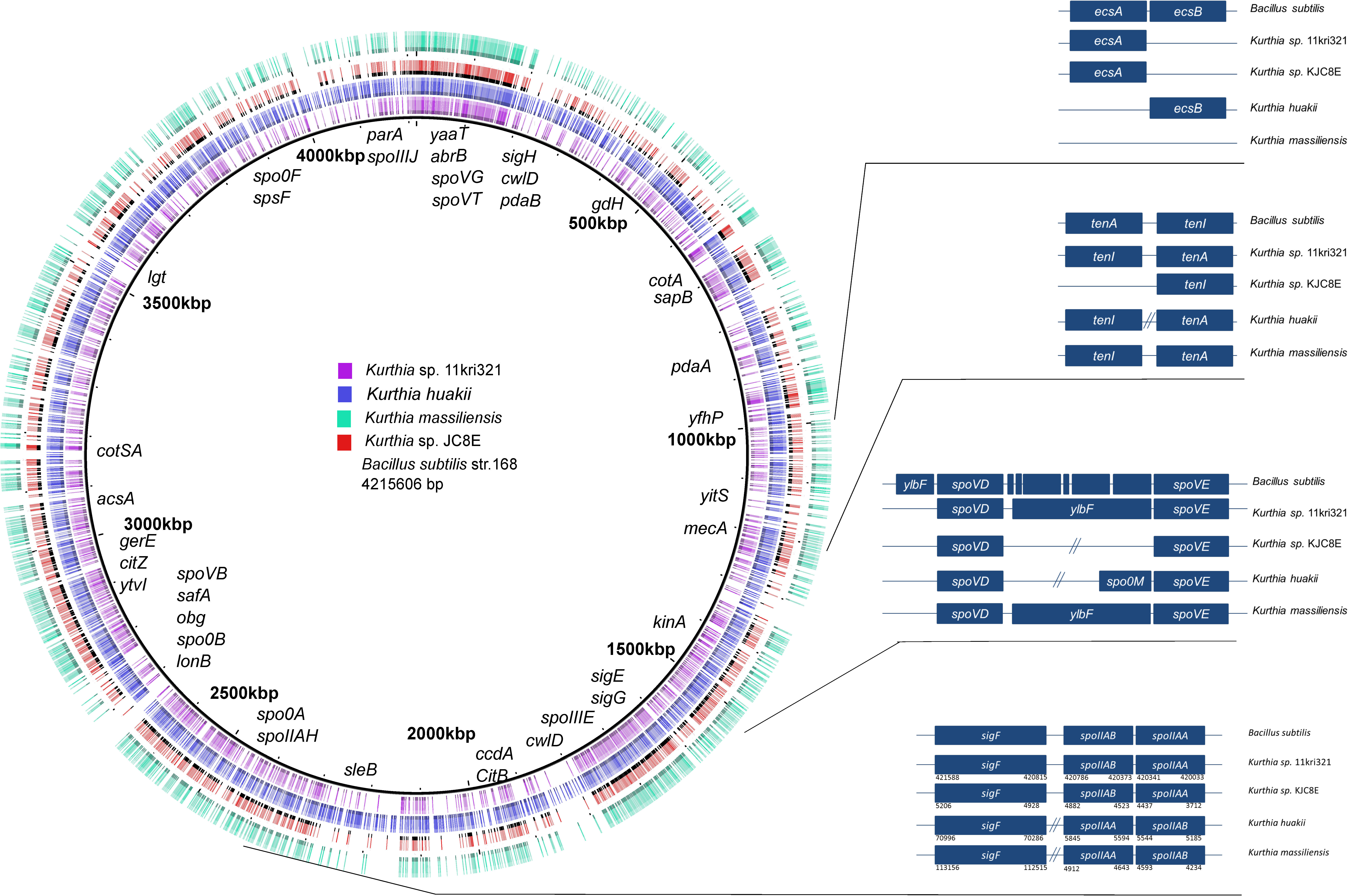
Comparison of the four *Kurthia* genomes publicly available to *Bacillus subtilis* subsp. *subtilis* str. 168. Genes related to sporulation are shown on the circular representation of the genomes. Four operons related to sporulation are also presented showing differences among the four *Kurthia* strains and *Bacillus subtilis* subsp. *subtilis* str. 168. The parallel lines in the operon representations show that in the respective strains, there is not an operon organization of these genes and that in contrary are distantly located in the chromosomes.

The process of sporulation in other *Kurthia* spp. was also investigated using 2-D cryo electron microscopy (cryo-EM). The results clearly demonstrated that an engulfment step and the production of a forespore within the mother cell can be observed during spore-formation in *Kurthia* spp. (Fig. 3B-C-D). Finally, mature spores in *K. massiliensis* showed a similar structure as those of *Kurthia* sp. str. 11kri321 (Fig. 5A-B), but the exosporium appeared to be more complex in *K. massiliensis* (Supplementary Fig. 3).

**Figure 5.**
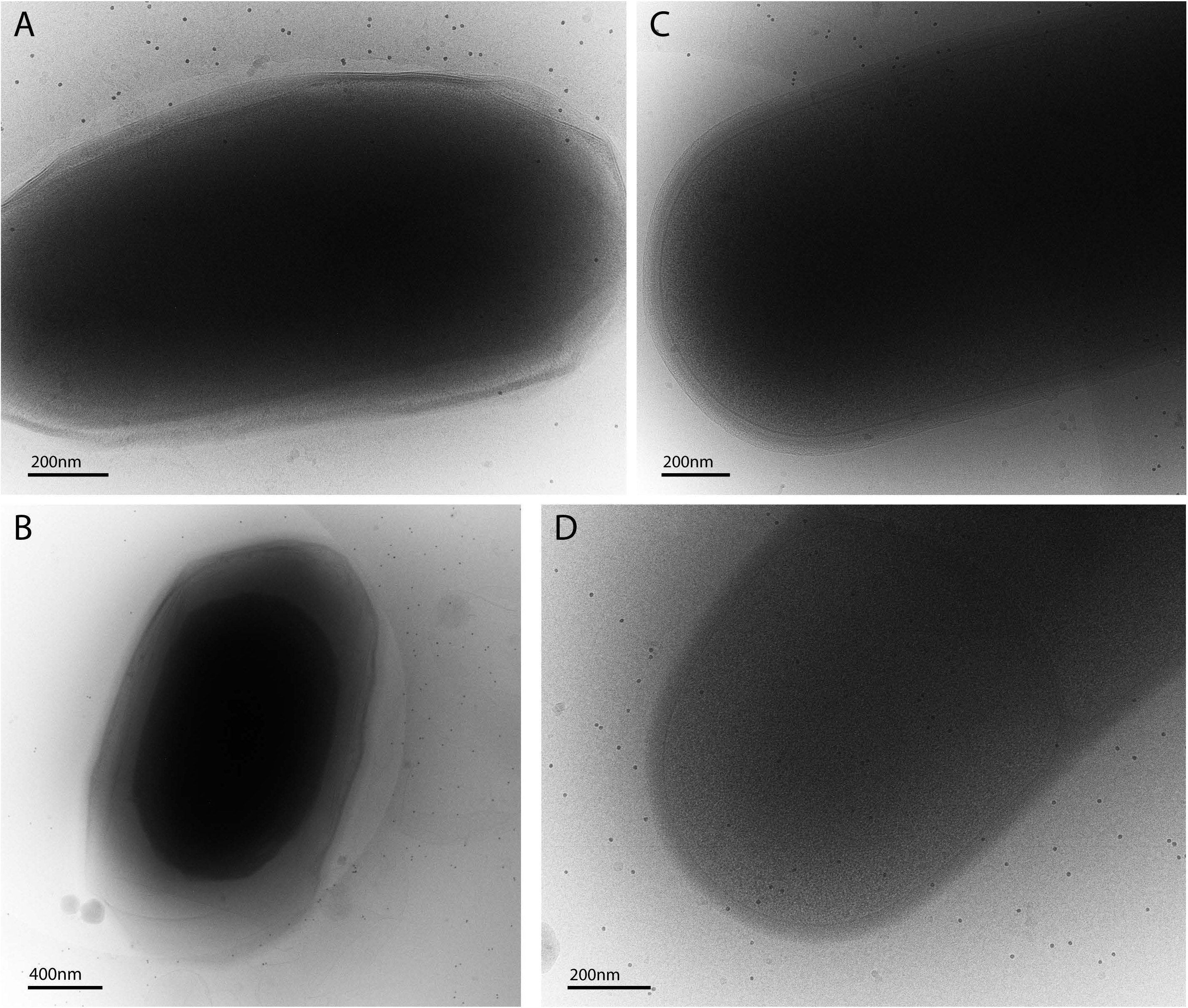
Two-D cryo-EM images of mature spores and vegetative cells of *Kurthia* spp. Spores of *Kurthia* sp. str. 11kri321 **(A)** and *K. massiliensis* str. JC30 **(B)** showed a similar structure and shape, however spore size is different: *Kurthia* sp. str. 11kri321 spores seem to be approximately double the size than those of *K. massiliensis* str. JC30. Because spores are constituted of concentrically arranged structures contributing to their high resistance (79), the external edges of spores look blurred compared to vegetative cells of *Kurthia* sp. str. 11kri321 **(C)** and *K. massiliensis* str. JC30 **(D)** in which the cytoplasmic membrane and cellular content are visible.

## Discussion

*Kurthia* is a genus described so far as encompassing asporogenic species. Although the conditions inside the sampled borehole were close to the optimal conditions for the vegetative growth of strain 11kri321, after nutrient starvation, we observed bodies that refracted light and retained malachite green staining (Fig. 1C and 1D), as it is the case for endospores produced by *B. subtilis*. Although, the number of sporulation events was too limited (Supplementary Fig. 2) for an in-depth characterization of the resistance of such bodies, the imaging data and germination study support the hypothesis that phase-bright bodies produced by *Kurthia* sp. str. 11kri321 correspond to spores (Fig. 2; Supplementary Movie 1).

The screening for known sporulation genes in the genome of *Kurthia* sp. str. 11kri321 and in other available genomes of representatives of the genus, suggests that sporulation in *Kurthia* spp. differs drastically from other Bacilli and deviates considerably from the proposed minimum core of sporulation genes (6). For instance, in *Kurthia* sp. str. 11kri321, only a small fraction of the genes (18 genes) belongs to those reported as conserved in all spore-forming Bacilli and Clostridia. Even a smaller number of the genes found in *Kurthia* sp. str. 11kri321 (14 genes), correspond to the 44 genes considered to be essential for sporulation in *B. subtilis* (6). A similar gene content was detected in the other *Kurthia* spp. investigated here, suggesting that the deviation to the sporulation pathway compared to other Bacilli is a feature conserved in the entire genus. The use of a comparative genomic approach to understand the evolution of endospore-formation has greatly contributed to our current knowledge of the distribution of endospore-formation genes within and outside Firmicutes (3, 6, 12). The presence of a conserved set of genes in endospore-forming Firmicutes with dramatically different lifestyles and even in endospore-forming Firmicutes with small and probably streamlined genomes, such as those of segmented filamentous bacteria (8, 9, 33), suggests that this set of conserved genes represents a close approximation to the true minimal set of sporulation genes (6). However, the discovery of sporulation in *Kurthia,* as well as the finding of a significant deviation in this genus from the accepted core of sporulation-related genes, extends our understanding of what constitutes a minimal set of sporulation-specific genes (34). Among the genetic elements involved in sporulation that are conserved in all Bacilli and Clostridia, the pathway responsible to enter in this energy-intensive process appear to be well preserved (17). Accordingly, the main regulators of this decision-making phase, SigH, the stationary phase sigma factor, as well as Spo0A, the master transcriptional regulator of sporulation, are highly conserved among endospore-forming Firmicutes (18, 21). This seems also to be the case in *Kurthia* spp. (Fig. 3A). Besides Spo0A, four highly conserved (24) major sigma factors (SigF, SigE, SigG and SigK) are essential for sporulation (35). All of the above were present in *Kurthia* spp. (with the exception of SigK in *K. senegalensis*), suggesting conservation in the directing elements responsible for differential gene expression in the mother cell and the forespore. This hypothesis is also supported by detection of homologs to SpoIIAA and SpoIIAB, two regulators of the expression of the sigma factors mentioned above (25, 26).

There are substantial differences in the sporulation process between Bacilli and Clostridia. One such difference is the way by which Spo0A is activated. In Clostridia, Spo0A is phosphorylated directly (36, 37), while in Bacilli the proteins Spo0B (phosphotransferase) and Spo0F (response regulator) are involved in a process known as phosphorelay (22). In the case of *Kurthia* sp. str. 11kri321 the presence of a homolog to *spo0F* and *obg* gene (Fig. 3) suggests that a phosphotransferase should be present. The same appears to be true for other members of the genus. In Bacilli, a family of sporulation-specific sensor histidine kinases (KinA, KinB, KinC, Kin D) is responsible for the activation of Spo0A (23), but not all members of this family are essential for signaling during sporulation (38). The detection of a gene encoding KinA in the genome of *Kurthia* sp. str. 11kri321 and *K. senegalensis* suggests a signaling pathway similar to that of other Bacilli. One of the most striking differences between the sporulation gene content in *Kurthia* spp. compared to other Firmicutes concerns the stage of engulfment. Engulfment is a remarkable example of a very rare phenomenon occurring in bacteria that allows the internalization of the cell membrane within the cytoplasm by a phagocytosis-like phenomenon (39). The so-called zipping process is a key mechanism operating during engulfment, but none of the genetic components identified so far in *Bacillus* spp. were detected in *Kurthia* sp. str. 11kri321 and the other species studied here. The process of engulfment normally requires three essential proteins: SpoIID, SpoIIM, and SpoIIP. The DMP complex plays a crucial role during septal thinning, which is the first step of the process of the degradation of septal peptidoglycan (39–41). From these three proteins, solely a gene encoding the peptidoglycan hydrolase SpoIID (40), was detected in the genome of *K. massiliensis*. A second mechanism for membrane migration mediated by SpoIIQ and SpoIIAH was also discovered in cells of *B. subtilis* in which peptidoglycan was removed enzymatically (39, 41). This suggests that more than one mechanism might ensure that engulfment is robust. Although none of the above could be detected in the genomic analysis of the *Kurthia* spp., the clear observation of the stages of membrane engulfment made by cryo-ET in the case of *K. huakuii* and *K. senegalensis* (Fig. 3B-C-D), as well as the conservation of differentially expressed sigma factors (for instance SigG and SigK) in most of the species (Fig. 3A), supports the conservation of endosporulation and indicates that an alternative engulfment mechanism should operate in the case of *Kurthia* spp.

The apparent absence of genes encoding for small acid-soluble proteins (SASPs) in the case of *Kurthia* spp. should also be discussed. In fact, these proteins are known to bind DNA and participate in its protection against heat, UV radiation and other damaging agents and represent up to 20% of the total spore proteins found in *B. subtilis* (42–45). However, the formation of viable spores does not require a great diversity of SASPs (46, 47), and this could support that no genes with homology to these SASPs were identified. Likewise, only a small number of genes involved in cortex formation were present in the analyzed genomes, therefore *Kurthia* spp. might have a different mechanism for the formation of this layer. In the same way, we identified a much smaller set of genes encoding proteins clearly related to the formation of the spore coat (only *cotA*; Fig. 3A). However, this might not be surprising as the distribution of spore coat proteins appears to reflect a differential adaptation of the organism to specific niches (6), and in consequence, are expected to be highly variable.

Dipicolinic acid (DPA), an important molecule in mature spores, has been used as a biomarker for detecting endospore-formers in environmental samples (31). However, none of the genes responsible for the production of DPA were detected in the genome of *Kurthia* sp. str. 11kri321 and the other species studied here. Previously it has been argued that unstable DPA-lacking spores fail to complete sporulation in *B. subtilis* (28, 29). However, it was also demonstrated that mutants lacking the *spoVF* operon could produce stable spores (not resistant to wet heat) under the condition that either *ger3* or *sleB* genes would also be absent (29, 30). These studies have shown that DPA does not influence the resistance of the spore to desiccation and dry heat, but yes to UV radiation (30). DPA-free spores tend to have lower core density but remain resistant and viable (48). As *Kurthia* sp. str. 11kri321 and the other *Kurthia* spp. lack the gene *ger3*, this suggests *Kurthia* spp. produces DPA-free stable spores better adapted to specific environmental insults. Moreover, an effect of increase in pH has been previously shown to lead to the release of DPA from *Alicyclobacillus acidoterrestris* spores (49). Therefore, it is likely that an adaptation to an extremely alkaline pH, such as the pH tolerated by *Kurthia* sp. str. 11kri321, has selected stable DPA-free spores in this species.

The overall results question the validity of describing organisms as asporogenic solely on the basis of the fact that sporulation has not been observed. Understanding the way in which endospore-formation, a seemingly ancestral characteristic, can be lost to give rise to truly asporogenic Firmicutes is essential to study the ecology and evolution of this clade. Considering that almost any natural habitat is subjected to significant variations in environmental parameters, it is difficult to conceive of a condition in which the complete loss of sporulation would be an advantage for Firmicutes. Accordingly, we propose to use the term *cryptosporulant* to designate those groups for which sporulation has not been investigated, and also for which a detailed physiological and genomic analysis is not yet possible to reliably define the capability of a clade to produce or not spores.

The discovery of sporulation in *Kurthia* sp. str. 11kri321 and other species of the genus paves the path for further investigation of cryptosporulant or asporogenic Firmicutes. The ecology of the few described species of *Kurthia* supports the cryptosporulant lifestyle of the genus. *Kurthia* strains have been isolated from diverse environments such as stool (50), biogas slurry (51), medical samples (50), cigarettes (52) and methanogenic bacterial complexes (53). In metagenomic studies, the genus *Kurthia* has been found in snail gastrointestinal tracts (54), restaurant kitchen cutting boards (55) and soy sauce fermentation processes (56, 57). In these habitats, sporulation can be a trait linked to survival and therefore a process that should be under selective pressure for conservation.

## Materials and Methods

### Sample collection and isolation

The geothermal reservoir of Krinides (N 41° 00.642’ E 024° 15.371’), near Philippoi, is situated in the Rhodope Massif (Kavala, Greece) (58). At the time of sampling, the water temperature at the output of the borepipe was 29.1°C, pH was 9 and conductivity 415 μS/cm. Water and biofilms from the outflow were collected in a sterile 1 L bottle and filtered through a 0.22 μm nitrocellulose membrane (Millipore, USA). The membrane was transported to the laboratory on ice and stored at 4 °C for bacterial enrichment into 10 mL of Nutrient Broth (Biolife, Italy). The enriched culture was then plated on Nutrient Agar (NA) and single colonies were obtained. Each colony was plated repeatedly to attain pure aerobic bacterial isolates. Colony morphology was observed after 12 h of growth. The capability to form spores was observed after starvation for 15 days using phase-contrast microscopy (Leica DM R, magnification 1000x). A differential staining for endospores and vegetative cells was performed using malachite green and safranine, as previously described (59). Gram-staining was also performed in order to determine the Gram character of the strain. Cell growth was monitored at different temperatures (4, 15, 20, 25, 35, 45, 50, 55, and 60°C) over 4 days in nutrient broth medium. To determine the pH range in which *Kurthia* sp. str. 11kri321 grows, nutrient broth medium at pH 4 to 13 was prepared (intervals of 0.5 pH unit), and growth was monitored at optimum growth temperature (25°C), over 4 days. All tests were performed in triplicates.

### Strain identification

#### gDNA extraction and sequencing

Genomic DNA was extracted from an overnight culture using the Genomic-tip 20/G kit (Qiagen GmbH, Germany). Sequencing was performed with the PacBio RS II system based on single molecule, real-time (SMRT) technology (Pacific Biosciences, California). The draft genome of *Kurthia* sp. str. 11kri321 presents a unique contig of 2964527 bases, and a G+C content of 36.7%. Genome annotation was performed using an Ergatis-based (60) workflow with minor manual curation and visualized with the Artemis Genome Browser and Annotation Tool (61). A total of 2893 coding sequences (CDSs), 82 tRNAs, and 27 rRNAs (9 copies of 16S, 23S and 5S rRNA genes) were predicted. This whole-genome project has been deposited at GenBank under the Bioproject PRJNA301103, and the Biosample ID SAMN04235798.

#### Induction of sporulation

Five sporulation media were prepared (SM1 with and without carbon source (32); SM2 with and without carbon source (31); Angle (62) with 10% glycerol). Pre-cultures of the four *Kurthia* strains (*K. massiliensis* str. JC30*, K. huakuii* str. LAM0618*, K. senegalensis* str. JC8E, and *Kurthia* sp. strain 11kri321) were inoculated overnight under optimal conditions. Biomass was retrieved with centrifugation at 6000 g for 3 min and transferred in the sterile sporulation media. The new cultures were incubated at optimal conditions for 7, 14, and 28 days. Presence of spores, vegetative, and dead cells was verified in the contrast-phase microscope and the three cell types were quantified in a Neubauer chamber.

#### Fluorescence and time-lapse microscopy

Cultures were grown either overnight (fresh cultures) or for 15 days (old cultures). The membrane dye FM4-64 (Invitrogen) was used at a final concentration of 500 ng/mL (diluted in DMSO) and incubated for 5 min before imaging. For time-lapse microscopy, a spore preparation (see below) was immobilized using a thin layer of Tryptic Soy Agar. Phase contrast microscopy images were taken at a sample frequency of 1 frame per 2 min. In both cases, images were acquired with an alpha Plan-Apochromatic * 100/1.46ph3 (Zeiss) on an Axio Imager M2 microscope (Zeiss) and a CoolSNAPHQ2 camera (Photometrics) controlled through Metamorph V7.5 (Universal imaging). Images were processed using ImageJ.

#### Electron cryotomography sample preparation and imaging

Cells grown on plates for 2 months were re-suspended in growth medium and frozen immediately for cryotomography studies. Mature spores and vegetative cells were collected from agar plates by resuspending them in growth medium and imaged with light microscopy at room temperature before and after freezing. In both cases, phase-bright objects were observed in the resuspension. Samples were then mixed with 20 nm colloidal gold particles, loaded onto glow-discharged carbon grids (R2/2, Quantifoil) and plunge-frozen into liquid ethane-propane mix cooled at liquid nitrogen temperatures with a FEI Mark IV Vitrobot maintained at room temperature and 70% humidity. The grids were imaged with a FEI Titan Krios TEM at 300 keV with a GIF2002 Imaging Filter (Gatan) and images were recorded on a 2Kx2K CCD Megascan model 795 camera (Gatan). Targets were picked randomly (n=35) and imaged. Images were acquired under low-dose conditions (final dose of 100 e^−^/Å^2^), 10 μm underfocus at 11500 × magnification, such that each pixel represented 7.9 Å. Tilt series were collected from −60° to +60° with 1 degree oscillation using EPU tomography software. Three-dimensional reconstructions were generated using the IMOD program (63).

Samples of cultures grown in liquid medium were treated as follows. 3.5 μl of vegetative cells or mature spores were mixed with 1 μl 10 nm Protein A-Gold (Department of Cell Biology of the University Medical Center, Utrecht, The Netherlands). 3.5 μl of that mixture was transferred to a 3 mm freshly glow-discharged Quantifoil R2/2 Cu 300 mesh holey carbon grid (Quantifoil Micro Tools GmbH, Jena, Germany). Excess liquid was blotted away using a Leica EM GP plunge-freezer at room temperature and 80% humidity (blot time 2-3 sec) and the grid was immediately plunge frozen in liquid ethane cooled by liquid nitrogen. Data was collected on a FEI Titan Krios TEM at 300 keV with a Quantum-LS energy filter (20 eV slit width) and a K2 Summit electron counting direct detection camera (Gatan).

Images of the cells and spores were recorded at magnifications of 1285 x, 7252 x and 11927 x, resulting in calibrated physical pixel size of 38.9 Å, 6.9 Å and 4.2 Å. The underfocus was changed between 100 μm and 7 μm with a total dose between 22 e^−^/Å^2^ and 0.7 e^−^/Å^2^. The images were recorded using the program SerialEM (64).

#### Phylogenetic analysis

16S rRNA gene sequences (>;1200 bp) of Firmicutes were retrieved from RDP (http://rdp.cme.msu.edu) (65) and aligned using the default parameters of MAFFT (66). A maximum likelihood phylogenetic tree was built using PhyML (67) and then graphics using the Newick utilities (68). Exhaustive HMM-based homology searches were carried out on a local genome databank of 253 Firmicute species by using the HMMER package (69) and as queries the HMM profiles of the complete set of 54 bacterial ribosomal proteins from the Pfam 29.0 database (http://pfam.xfam.org (70)). Twelve ribosomal proteins were discarded due to their absence from >50% of the considered genomes and because they contained paralogous copies making identification of orthologues difficult.

The remaining 42 single protein data sets were aligned with MAFFT v7.222 (66) with the L-INS-I algorithm and default parameters, and unambiguously aligned positions were selected with BMGE 1.1 (71) and the BLOSUM30 substitution matrix. Single protein datasets were concatenated by allowing a maximum of 4 missing taxa, resulting in a character supermatrix containing 5254 amino acid positions. A Bayesian tree was calculated from the ribosomal protein concatenate with PhyloBayes v3.3 (72) and the evolutionary model GTR+CAT+G4 (73). Two independent chains were run until convergence, assessed by evaluating the discrepancy of bipartition frequencies between independent runs. The first 25% of trees were discarded as burn-in and the posterior consensus was computed by selecting one tree out of every two. The tree was rooted based on previous evidence (2). The tree was visualized and metadata was added to it using iTOL (74).

#### Retrieval of sporulation genes sequences

Complete and draft sequences of spore-forming Firmicutes were downloaded from Comprehensive Microbial Resource (CMR) and Integrated Microbial Genome (IMG) websites. Search for spore-related genes was based on gene function category sporulation (CMR; sporulating category in IMG). The CMR version was 24.0 data release and the IMG version was 3.0. In addition to the protein sequence, nucleotide sequences including a 50-bp flanking region at both 5’- and 3’-ends were downloaded. Additional information on all retrieved genomes was obtained from the GenBank database.

#### Sequence data analysis

The *Kurthia* sp. Str. 11kri321 genome sequence was scanned for orthologs of the Firmicute core sporulation genes (as protein sequences) with TBLASTN (75), using default parameters and an E-value cutoff of 1E-11. A TBLASTN run on the shuffled protein sequences as a negative control set showed no hit with an E-value lower than 4e-4. The hits were ordered by position on the *Kurthia* sp. Str.11kri321 genome and inspected manually. The above procedure did not detect orthologs of SpoVFA and SpoVFB, therefore we attempted to detect those by pairwise dynamic-programming alignment. The protein sequences of SpoVFA and SpoVFB were each compared to all sequences of the *Kurthia* proteome using Needleman and Wunsch’s algorithm (76), as implemented by EMBOSS’s `needle` program (77) No hits were found. The publicly available genomes of *Kurthia huakuii* str. LAM0618, *Kurthia massiliensis* str. JC30, and *Kurthia senegalensis* str. JC8E were scanned for sporulation gene orthologs as described above. The comparison of the four *Kurthia* genomes to *Bacillus subtilis* subsp. *Subtilis* str. 168 was plotted using BRIG (78), followed by further manually curation.

The sequences of the *Kurthia* genomes analyzed herein were retrieved from GenBank under accession numbers: *Kurthia huakuii* str. LAM0618: NZ AYTB00000000.1, *Kurthia massiliensis* str. JC30: NZ CAEU00000000.1, and *Kurthia senegalensis* str. JC8E: NZ CAEW00000000.1. The genome of *Bacillus subtilis* subsp. *Subtilis* str. 168: NC_000964.3 was also retrieved.

#### DPA measurement

The presence of dipicolinic acid (DPA) in the spores was assessed, according to a previously published method (31). Fluorescence was measured with a Perkin-Elmer LS50B fluorometer. The excitation wavelength was set at 272 nm with a slit width of 2.5 nm. Emission was measured at 545 nm (slit width 2.5 nm). The device was set in the phosphorescence mode (equivalent to time-resolved fluorescence). The delay between emission and measurement was set at 50 μs. Measurements were performed every 20 ms. The integration of the signal was performed over a duration of 1.2 ms. Values recovered for each measurement corresponded to the mean of the relative fluorescence unit (RFU) values given by the instrument within the 30 s following sample introduction in the device. Finally, to transform RFU units into DPA concentrations, a 10-point standard curve was established using increasing concentrations of DPA from 0.5 μm up to 10 μm.

## Acknowledgments

We acknowledge funding from the Swiss National Science Foundation projects 31003A_132358/1, 31003A_152972, 31003A_179297 (PJ), 31003A_162716 (PHV), and 31003A_17002 (TBB), 179520 (BZ) from Fondation Pierre Mercier pour la science (PJ), and from REGARD for equality of women in science (SF). Further, we would like to acknowledge the Natural Sciences and Engineering Research Council of Canada for funding E.I.T. We acknowledge bioinformatics support by the de.NBI network, BMBF grant FKZ 031A533 to J.B. E.I.T would like to acknowledge NSERC (Natural Sciences and Engineering Research Council of Canada) for cryotomography data collection at the Facility for Electron Microscopy Research of McGill University and Dr. Kaustuv Basu for his help with data collection. We would like to thank Mrs. Nicole Jeanneret for her assistance in the isolation and characterization of the strain, Dr. Guillaume Cailleau for the 3D representation of the sporulation and germination models, and Prof. Dr. Henning Stahlberg for the support and use of the electron microscope. Instrumentation supported by the Microscopy Imaging Center (MIC) of the University of Bern was used for this study. Individuals responsible for funding this work did not participate in study design, data collection and interpretation, or the decision to submit this work for publication. The authors declare no conflict of interest. Any genetic information downloaded from Genbank should be considered to be part of the genetic patrimony of Greece, the country from which the sample was obtained. Users of this information agree to 1) acknowledge Greece as the country of origin in any country where the genetic information is presented and 2) contact the CBD focal point and the ABS focal point identified in the CBD website http://www.cbd.int/information/nfp.shtml if they intend to use the genetic information for commercial purposes.

## Supplementary Material

**Supplementary Figure 1. DPA measurement in spores of *Kurthia* sp. str. 11kri321 and in positive/negative controls.** Measurements have been performed in 5 replicates in *Kurthia* sp. str. 11kri321 spore preparations, in *Bacillus subtilis* spore preparations (positive control) and in *Serratia marcescens* vegetative cells (negative control). (*Kurthia* sp. str. 11kri321: mean DPA concentration (mDPA)= 0.779 μM, standard deviation (SD)= 0.00645. *B. subtilis*: mDPA= 31.0 μM, SD= 0.0345. *S. marcescens*: mDPA= 0.16 μM, SD= 0.0154).

**Supplementary Figure 2. Frequency of sporulation on different culture media for four *Kurthia* strains. (A)** All *Kurthia* strains were able to produce phase-bright bodies on most of the sporulation media tested **(B)**. After 7 days, frequency of spores, compared to vegetative cells and dead ones, never exceeded 10% (SM1C= Sporulation medium 1 with carbon source; SM1N= Sporulation medium 1 without carbon source; SM2C= Sporulation medium 2 with carbon source; SM2N= Sporulation medium 2 without carbon source; GLY= Angle medium with 10% glycerol).

**Supplementary Figure 3. Two-D cryo-EM images of spores of *Kurthia massiliensis*. (A-B)** Spores of *K. massiliensis* str. JC30 showed a complex exosporium with potentially modular protein structures (arrows).

**Supplementary Movie 1. Germination of *Kurthia* sp. str. 11kri321.** From a spore preparation immobilized on a thin layer of Tryptic Soy Agar, time-lapse microscopy showed transition from spores (phase-bright) to vegetative cells (phase-dark). Phase contrast microscopy images were acquired at a frequency of 1 frame per 2min.

**Supplementary Table 1. Sporulation genes of endospore-forming Firmicutes.** This table summarizes the presence or absence of homologs for 213 sporulation genes in 34 endospore-forming Firmicutes.

